# Genomic Surveillance of SARS CoV2 in COVID-19 vaccinated healthcare workers in Lebanon

**DOI:** 10.1101/2022.06.06.494965

**Authors:** Habib Al Kalamouni, Farouk F. Abou Hassan, Mirna Bou Hamdan, Andrew J. Page, Martin Lott, Nada Ghosn, Alissar Rady, Rami Mahfouz, George F. Araj, Ghassan Dbaibo, Hassan Zaraket, Nada M. Melhem, Ghassan M. Matar

## Abstract

The emergence of SARS-CoV-2 variants including the Delta and Omicron along with waning of vaccine-induced immunity over time contributed to increased rates of breakthrough infection specifically among healthcare workers (HCWs). SARS-CoV-2 genomic surveillance is an important tool for timely detection and characterization of circulating variants as well as monitoring the emergence of new strains. Our study is the first national SARS-CoV-2 genomic surveillance among HCWs in Lebanon. We collected 250 samples from five hospitals across Lebanon between December 2021 and January 2022. We extracted viral RNA and performed whole genome sequencing using the Illumina NextSeq 500 platform. A total of 133 (57.1%) samples belonging to the Omicron (BA.1.1) sub-lineage were identified, as well as 44 (18.9%) samples belonging to the BA.1 sub-lineage, 28 (12%) belonging to the BA.2 sub-lineage, and only 15 (6.6%) samples belonging to the Delta variant sub-lineage B.1.617.2. These results show that Lebanon followed the global trend in terms of circulating SARS-CoV-2 variants with Delta rapidly replaced by the Omicron variant. This study underscores the importance of continuous genomic surveillance programs in Lebanon for the timely detection and characterization of circulating variants. The latter is critical to guide public health policy making and to timely implement public health interventions.

## Introduction

Since its emergence in December 2019, severe acute respiratory syndrome coronavirus 2 (SARS-CoV-2) remains a global public health threat. As of May 16, 2022, more than 521,476,365 confirmed cases and 6,263,965 deaths have been reported worldwide [1]. In Lebanon, more than 1,098,030 confirmed cases and 10,408 deaths have been reported as of May 15, 2022 [2]. Vaccine development against SARS-CoV-2 proceeded in an unprecedented pace with 11 vaccines granted emergency use listing (EUL) by the World Health Organization (WHO) as of May 20, 2022 [3]. More than 11.4 billion vaccine doses have been administered worldwide since the start of COVID-19 vaccine rollout in December 2020 [1]. Despite the availability of effective vaccines against SARS-CoV-2, reports of breakthrough infections among vaccinated individuals are increasingly reported globally [4-8].

Since the emergence of SARS-CoV-2, new variants have evolved from the original SARS-CoV-2 strain (Wuhan 19 strain (WA1/2020)). These variants are classified into variants under monitoring (VUM), variants of interest (VOI), and variants of concern (VOC) and [9, 10]. VUM are variants with genetic changes that are suspected to affect virus characteristics and may pose future risk but with yet no clear evidence of phenotypic or epidemiological impact. VOI are SARS-CoV-2 variants possessing predicted or known genetic changes that affect the characteristics of the virus (transmissibility, disease severity, immune escape, therapeutic escape) and known to cause significant community transmission or multiple COVID-19 clusters in multiple countries. VOC are SARS-CoV-2 variants that meet the definition of a VOI and are associated with increased transmissibility detrimental change in COVID-19 epidemiology, increased virulence, changed clinical disease presentation, and decreased effectiveness of public health and social measures, vaccines, or therapeutics against the virus. To date, the World Health Organization (WHO) has identified five VOCs worldwide: Alpha (B.1.1.7 lineage) first detected in the United Kingdom (UK), Beta (B.1.351 lineage) first detected South Africa, Gamma (P.1 lineage) first detected in Brazil, Delta (B.1.617.2 lineage) first detected in India and Omicron (B.1.1.529 lineage) first reported in South Africa [10]. Moreover, several VOI have been identified in several countries; these include B.1.427 and B.1.429 from the USA (California, WHO alert since July 6, 2021), B.1.525 from the United Kingdom and Nigeria, B.1.526 from the USA (New York), B.1.617.1 and B.1.617.3 from India, P2 from Brazil, and C.37 from Peru [9]. The WHO is continuously monitoring and assessing the evolution of SARS-CoV-2 and the emergence of new variants with increased risk to the global public health.

The B.1.617.2 lineage along with its sublineages made up the Delta variant that was responsible for the COVID-19 surge in India, eventually spreading and dominating globally [11]. Despite the high replicative efficiency, reduced sensitivity to host immune responses, and high transmissibility of the Delta variant compared to previous VOCs, vaccine effectiveness was sustained against both the Alpha and Delta variants [11]. In November 2021, the surge of cases in South Africa marked the identification of a new VOC named Omicron (B.1.1529). Omicron replaced the Delta variant and was characterized by a higher number of amino acid substitutions, higher transmissibility and partial resistance to vaccine induced immunity compared to previous VOCs [12-14]. Studies showed that although Omicron had higher rates of reinfection, it was clinically less severe compared to the Delta variant suggested to be driven by prior infections and T cell immune responses [11]. While two doses of COVID-19 vaccines elicit high level of protection against symptomatic disease, the former wanes 4-6 months following the second dose of the BNT162b2 (Pfizer-BioNTech), mRNA-1273 (Moderna) or ChAdOx1 nCoV-19 (Oxford-AstraZeneca vaccines) [15, 16]. Recent studies showed that vaccine effectiveness against the Omicron variant (B.1.1.529) was lower than the Delta variant (B.1.617.2) after primary immunization with 2 doses of the ChAdOx1 nCoV-19, BNT162b2 or mRNA-1273 vaccines with significant reduction in vaccine effectiveness against the two variants ≥25 weeks following the second dose [17]. Nevertheless, vaccine effectiveness against symptomatic disease was restored following a booster shot, underscoring the importance of a third dose [15, 18].

Healthcare workers (HCWs) are at increased risk of SARS-CoV-2 infection compared to the general population and have been prioritized in COVID-19 vaccine rollout worldwide [19]. In Lebanon, HCWs were among the first priority groups to receive the BNT162b2 vaccine; the administration of the latter started in mid-February 2021 and the second dose was administered 21 days following the first dose. The use of personal protective equipment (PPE) including mask mandates for patients and visitors significantly reduced the occupational risk of acquiring COVID-19 by HCWs [20]. However, waning immunity and the emergence of antigenically drifted VOCs meant that HCWs are at risk of breakthrough infections [4, 5, 8, 21]. Among 22,729 HCWs in the US who received at least one dose of an mRNA-based vaccine (BNT162b2 or mRNA-1273), 189 tested positive for SARS-CoV-2 [6]. While the majority (60%) of these infections occurred within 14 days following the first dose, 14% of the cases occurred >14 days post second dose. Similarly, 39 out of 1,497 HCWs in Israel had breakthrough infections after receiving the second dose of BNT162b2 vaccine with most of these infections being asymptomatic or mild [4]. Moreover, the incidence of breakthrough infections following the second dose of the ChAdOx1 nCoV-19 vaccine was estimated at 1.6% among 3000 HCWs in India [7]. Interestingly, the Omicron variant requires 20 to 40 times more neutralization antibodies than Delta, which might have contributed to higher breakthrough infection among people with only two vaccine doses [22]. The resurgence of SARS-CoV-2 is attributed to waning of immunity over time and emergence of SARS-CoV-2 variants [23]. These breakthrough infections carry an infectious potential especially since these infections are mostly asymptomatic and thus would increase the risk of viral spread to high-at-risk populations [4].

Genomic surveillance of SARS-CoV-2 was first initiated in Lebanon back in 2020 with direct support from the WHO [24]. Lebanon was the first in the Eastern Mediterranean Region (EMR) to identify the Delta variant in a timely manner. However, the sampling was done randomly and mainly on travelers. In collaboration with the Ministry of Public Health (MoPH) and direct support from the WHO, this study allowed for the establishment of a structured mechanism by which we can perform genomic surveillance of SARS-CoV-2 on HCW’s to support the timely implementation of public health measures to control the spread and emergence of new SARS-CoV-2 variants and inform global surveillance. Here, we report the first genomic data from this surveillance effort focusing primarily on breakthrough infections detected among HCWs, a group with the highest vaccination coverage among the Lebanese population.

## Results

### Demographic characteristics of study participants

Our study included a total of 250 HCWs testing positive by RT-PCR between December 1, 2021 (n=27), and January 31, 2022 (n=223). Data on the occupation and COVID-19 vaccine status were available for only 175 HCWs working at AUBMC. Among those, the majority were females (64%) and received at least two doses of the BNT162b2 COVID-19 vaccine (95.4%). Nurses (39.4%), medical doctors (14.8%) and technicians (10.3%) accounted for the majority of samples (Table 1). None of the SARS-CoV-2 positive HCWs were hospitalized.

**Table 1.** Demographic characteristics of HCWs recruited at AUBMC.

| Variable | N | % |
| --- | --- | --- |
| **Gender** |  |  |
| Males | 63 | 36 |
| Females | 112 | 64 |
| **Vaccination status** |  |  |
| ≥2 doses | 167 | 95.4 |
| 1 dose | 4 | 2.3 |
| Unvaccinated | 1 | 0.6 |
| Not documented | 3 | 1.7 |
| **Occupation** |  |  |
| Nurses | 69 | 39.4 |
| Medical doctors | 26 | 14.8 |
| Technicians | 18 | 10.3 |
| Clerks, tellers and clinic assistants | 16 | 9.1 |
| Pharmacists | 5 | 2.8 |
| Physical therapists | 2 | 1.1 |
| Others<sup>a</sup> | 40 | 22.8 |

### SARS-CoV-2 phylogenetic analysis

Lineage analysis was performed using the Pangolin COVID-19 lineage assigner. Overall, 10 lineages were identified among HCWs (Table 2). A total of 17 samples did not yield sufficient sequencing data to provide a lineage. This was most likely due to sample storage and handling errors as they passed through multiple labs in multiple countries. Consequently, we excluded them from the analysis. As expected, the Omicron variant was the predominant VOC (90.6%) detected in most of our analyzed samples followed by the Delta variant (6.4%). The predominant lineages identified were BA.1.1 and BA.1 accounting for 57.1% and 18.9% of the samples, respectively (Table 2 and Fig 1). Collection date was available for 225 of the specimens. Out of those, 27 were collected in December 2021 and 198 were collected in January 2022. Our results revealed that Omicron BA.1.1 variant was the predominant VOC circulating in December 2021 (37%) and January 2022 (56%) followed by 22.2% and 19.2% BA.1, respectively (Fig 2).

**Table 2.**
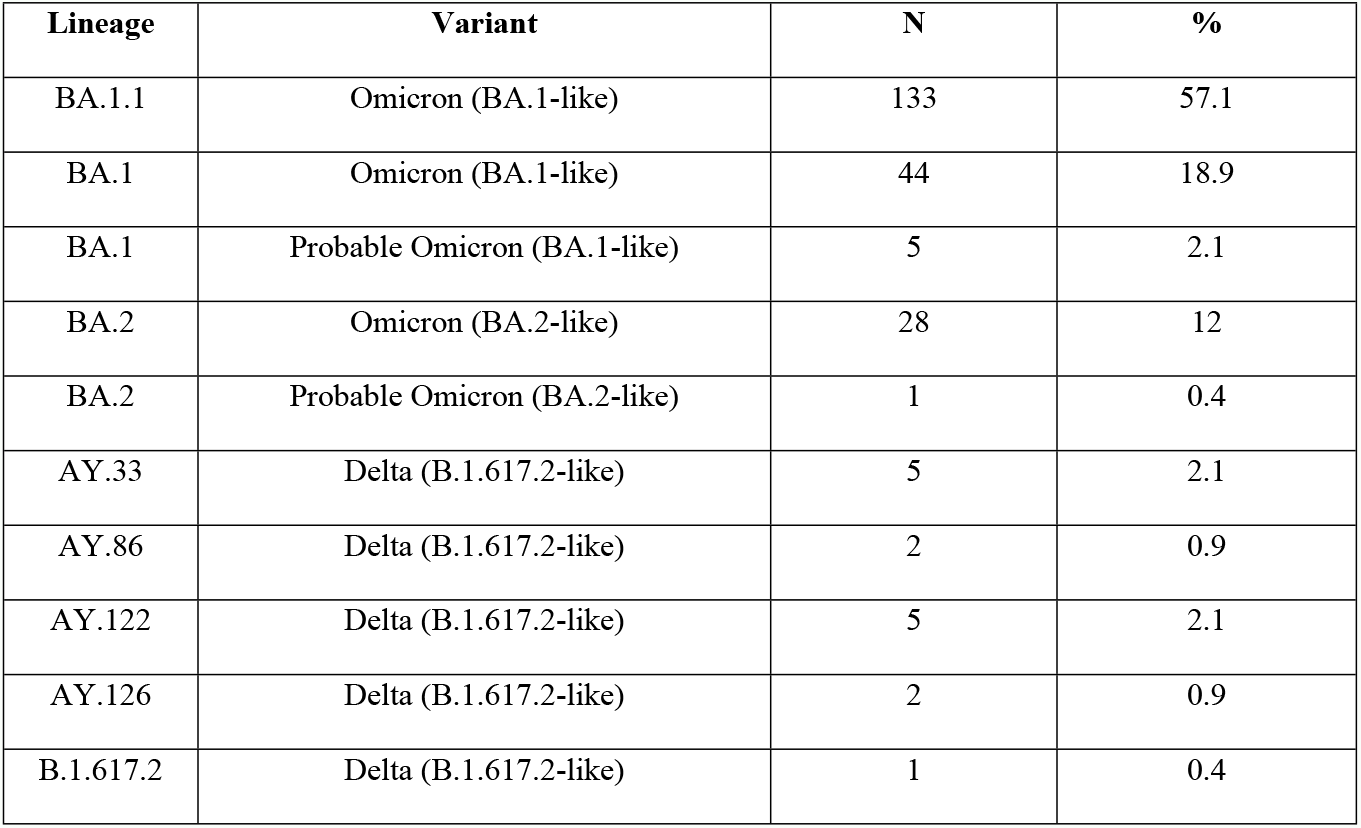
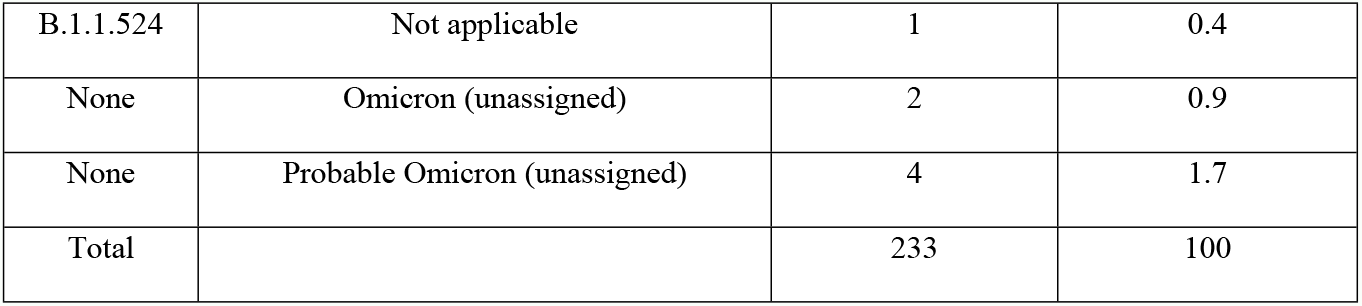
SARS-CoV-2 lineages and variants detected in HCWs.

| Lineage | Variant | N | % |
| --- | --- | --- | --- |
| BA.1.1 | Omicron (BA.1-like) | 133 | 57.1 |
| BA.1 | Omicron (BA.1-like) | 44 | 18.9 |
| BA.1 | Probable Omicron (BA.1-like) | 5 | 2.1 |
| BA.2 | Omicron (BA.2-like) | 28 | 12 |
| BA.2 | Probable Omicron (BA.2-like) | 1 | 0.4 |
| AY.33 | Delta (B.1.617.2-like) | 5 | 2.1 |
| AY.86 | Delta (B.1.617.2-like) | 2 | 0.9 |
| AY.122 | Delta (B.1.617.2-like) | 5 | 2.1 |
| AY.126 | Delta (B.1.617.2-like) | 2 | 0.9 |
| B.1.617.2 | Delta (B.1.617.2-like) | 1 | 0.4 |
| B.1.1.524 | Not applicable | 1 | 0.4 |
| None | Omicron (unassigned) | 2 | 0.9 |
| None | Probable Omicron (unassigned) | 4 | 1.7 |
| Total |  | 233 | 100 |

**Fig 1.**
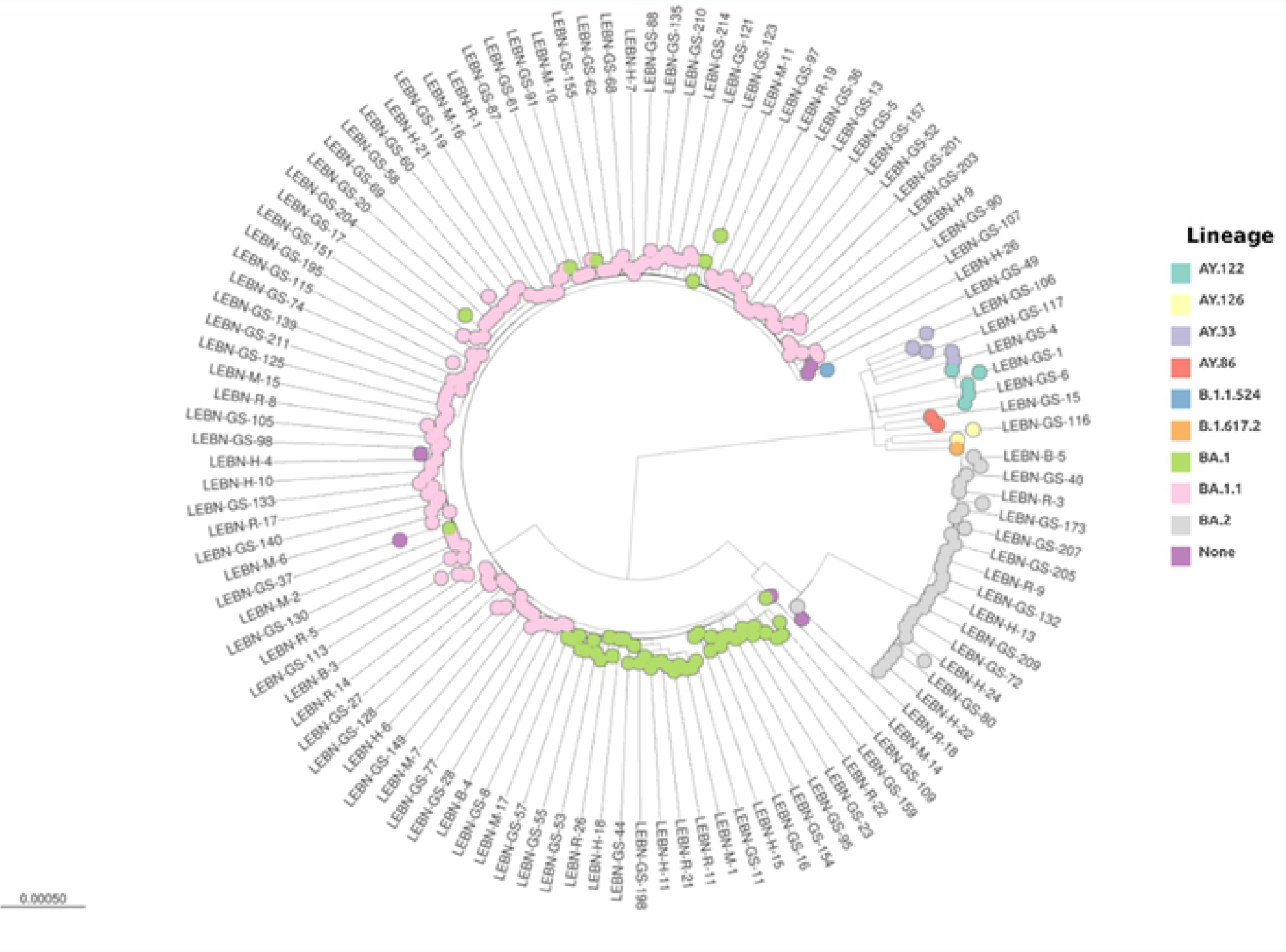
Phylogenetic analysis. Phylogenetic analysis of SARS-CoV-2 genome in 250 samples collected from HCWs in Lebanon. Each lineage is specified with a unique color.

**Fig 2.**
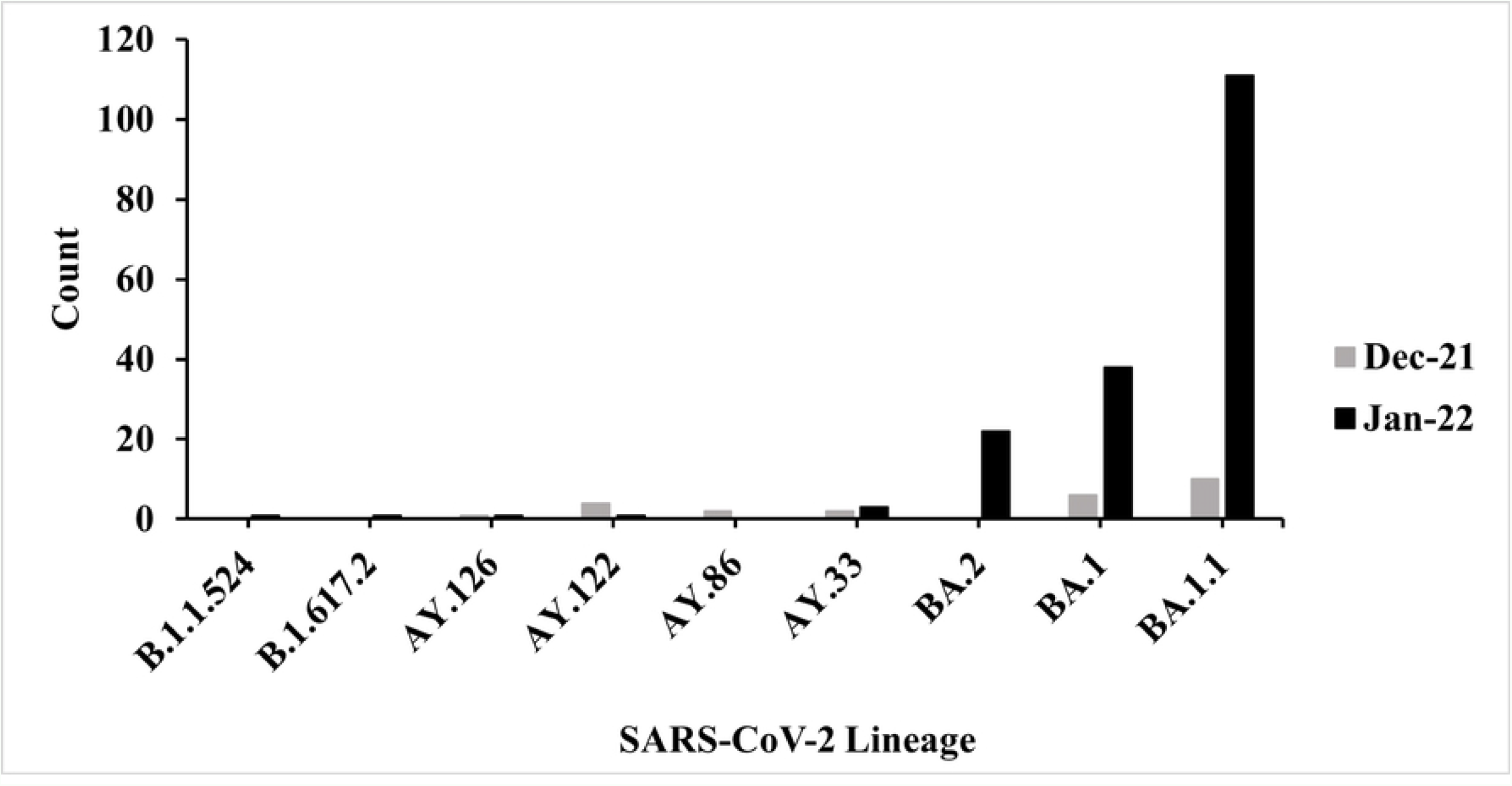
Frequency of SARS-CoV-2 lineages among HCWs. Data present the number of samples with a specific SARS-CoV-2 lineage detected in December 2021 (n=27) and January 2022 (n=198) out of 225 samples with available date of PCR testing.

## Discussion

Whole genome sequencing is important to characterize circulating SARS-CoV-2 variants and to detect emerging variants. In Lebanon, data on SARS-CoV-2 genome sequencing are lacking specifically among HCWs who are at high risk of acquiring the infection. Genomic analysis of 11 specimens collected early during the pandemic in Lebanon (February– March 2020) showed that the B.1 lineage was the most prominent, followed by the B.4 lineage and the B.1.1 lineage [33]. Between February 2020 and January 2021, the most frequently reported SARS-CoV-2 lineage among 58 analyzed samples was B.1.398 followed by B.1.1.7 and B.1 [34]. Moreover, an analysis of 905 samples showed the rapid emergence and dominance of the B.1.1.7 Alpha variant between January and April 2021 followed by the replacement of Alpha with Delta variant between June and July [24]. Our study reveals that Omicron BA.1.1 followed by BA.1 predominated during January 2022.

HCWs were identified as a high-priority group for COVID-19 vaccination by the WHO Strategic Advisory Group of Experts framework for the allocation and prioritization of COVID-19 vaccination [35] and the Advisory Committee on Immunization Practices [36]. Consequently, many countries including Lebanon designated HCWs as a priority group for vaccination [37]. Despite the rollout of effective COVID-19 vaccines, breakthrough infections have been increasingly reported worldwide specifically among HCWs [4, 8, 21, 38]. The incidence of breakthrough infections among vaccinated HCWs is low and it was recently estimated at 0.011 to 0.001 per 100 individuals [39]. Nevertheless, these infections pose a risk of transmission to vulnerable populations as most of these breakthrough infections are mild or asymptomatic [4, 39]. In addition to waning of vaccine-induced immunity over time, genetic variants of SARS-CoV-2 also affect vaccine-induced immune responses [23]. Compared to Alpha and Delta variants, the Omicron variant causes higher rates of breakthrough infection and lower hospitalization rates [23, 40]. Moreover, the transmissibility of Omicron is higher than its predecessors; this is mainly attributed to the high number of mutations in the spike protein [12, 41]. The high number of mutations contributed to 3-fold higher binding affinity of the RBD of Omicron to the ACE2 receptor compared to Wuhan HU-1 and Delta [42, 43]. Studies also showed that vaccine effectiveness against Omicron variant was lower than that of Delta and that neutralizing antibody activity against Omicron is significantly lower than Beta and Delta variants [17, 23, 44].

In the Middle East and North Africa (MENA) region, there are scarcity of data on vaccine effectiveness against emerging SARS-CoV-2 variants. There are also limited data on breakthrough infections among HCWs following vaccination. Multiple studies in Qatar reported on vaccine effectiveness and breakthrough infections following vaccination [45-48]. The estimated cumulative incidence of breakthrough infections in Qatar was at 0.59% after a median of 89 days from receiving the second dose of the mRNA-1273 vaccine and 0.84% after receiving the BNT162b2 vaccine [45]; waning of vaccine effectiveness against SARS-CoV-2 infection was also reported at 4 months following the second dose of the BNT162b2 vaccine [47]. Moreover, mRNA-1273 vaccine effectiveness against symptomatic infection caused by the B.1.1.7 or B.1.351 variants in Qatar was estimated at 98.6% ≥14 days after receiving the second dose [48].

The Omicron variant (B1.1.529) was first detected in early November 2021 in multiple countries and has been designated as a VOC by the WHO on November 26, 2021 [49]. Four Omicron lineages were first identified: B1.1.529, BA.1, BA.2, and BA.3 [50]. Recently, four novel Omicron subvariants designated as BA.4, BA.5, BA.2.12.2 and BA.2.13 have also emerged and started spreading globally [51-54]. While BA.4 and BA.5 account for 50% of new sequenced samples in South Africa, BA.2.12.1 and BA.2.13 account for nearly 30% and 5% of new cases in the United Sates and Belgium, respectively [51]. Compared to BA.2, these novel omicron subvariants exhibit additional mutations in the spike region namely L452Q for BA.2.12.1, L452M for BA.2.13, and L452R+F486V for BA.4 and BA.5 [51]. These mutations have been shown to provide potential immune escape characteristics and higher transmission than BA.2 [51, 53]. In this study, we found that Omicron BA.1.1 and BA.1 lineages were the predominant circulating lineages in our cohort of HCWs between December 2021 and January 2022. Delta variant was detected in only 6% of our samples suggesting the replacement of the Delta variant with Omicron as the predominant circulating VOC which is consistent with global trends observed during the same period [55]. We did not detect BA.3 lineage or any recombinant lineages in our sequenced samples. The former does not have specific mutations in the spike protein but rather a combination of mutations from BA.1 and BA.2 [56]. The rate of spread of the three Omicron lineages (BA.1, BA.2 and BA.3) differs with BA.1 and BA.2 being the predominant lineages. Between December 2021 and January 2022, BA.1 lineage accounted for 78% of sequenced samples submitted to the GISAID database compared to 16% of BA.2 [57]. This is similar to our findings reflecting the predominance of BA.1 over BA.2.

The subvariant BA.2 shares 32 mutations with BA.1 but has distinct 28 mutations, four unique ones in the RBD region alone, which according to a deep learning algorithm, made it far more likely than other lineages to be the next dominant subvariant [50]. Indeed, BA.2 had already become the dominant variant in multiple countries such as Denmark and UK in February 2022 [58]. Moreover, as of May 16, 2022, 78% of sequenced samples submitted to GISAID database were BA.2 compared to 5% BA.1. BA.2.12.1 and BA.4 accounted for 13% and 3% of submitted sequences, respectively. As we continue our national genomic surveillance beyond January 2022, we expect a shift in dominance in favor of the highly transmissible BA.2 subvariant and as well as an expected detection of other Omicron subvariants.

Our study has several limitations. Our study did not include samples from all regions in Lebanon and thus is not fully representative of the situation in Lebanon. However, given that Beirut is the capital of this small country and sees a lot of population movement during the week, and particularly on the weekends when its residents travel to their villages across Lebanon, we believe that our data from Beirut are to some extent representative of the whole country. We were also unable to gather clinical data of HCWs, which hampered our ability to assess risk factors associated with Omicron breakthrough infections. Moreover, data on receiving the date of the second and booster shots were unavailable and thus we were unable to estimate vaccine effectiveness between the date of receiving the booster and the date of breakthrough infection.

## Conclusion

Our findings underscore the importance of continuing genomic surveillance in Lebanon in order to monitor virus evolution and the emergence of novel SARS-CoV-2 variants. This is particularly important in HCWs as they are more likely to be exposed to emerging variants and can act as an advanced warning proxy to the wider community. More recently, two Omicron lineages (BA.4 and BA.5) have been identified in South Africa before being detected in several countries worldwide including Botswana, Belgium, Denmark, the United Kingdom, France, Germany, Portugal and China [54, 59, 60]. Therefore, continuing genomic surveillance will help assessing the characteristics and the public health implications of these lineages and other variants that might emerge and contribute to more informed public health intervention strategies.

## Materials and Methods

### Study design, population, and data collection

This study is part of a national surveillance program in collaboration with the Epidemiological Surveillance Unit (ESU) at the Lebanese MoPH. Accordingly, a waiver of informed consent was granted by the Institutional Review Board (IRB) at the American University of Beirut (AUB). Between December 1, 2021, and January 31, 2022, nasopharyngeal swabs were collected from a total of 250 COVID-19-positive HCWs from five Lebanese healthcare centers. Samples with *Ct values* of less or equal to 25 were used. The majority of samples (n=205) were collected from three hospitals in Beirut: AUB Medical Center (n=175), Rasoul Al Aazam Hospital (n=25) and Belle Vue Hospital (n=5). The remaining were collected from Hammoud Hospital in South Lebanon (n=26) and Mount Lebanon Hospital in Mount Lebanon (n=19). Aliquots of the collected samples were stored at -80°C until processed. The date of positive PCR, vaccination status, specific occupation, and hospitalization status of participants were collected.

### RNA extraction and whole genome sequencing (WGS)

Aliquots of the nasopharyngeal swabs (140 µl) were used to extract total RNA following manufacturer’s instructions (QIAamp Viral RNA mini-Kit, QIAGEN, Hilden, Germany, Cat. 52906). Aliquots were eluted in 30 µL of Buffer AVE. Both the concentration and quality of RNA samples were measured and checked with the Denovix Blue DS-11 Spectrophotometer. Viral RNA extracts were sequenced at the Quadram Institute Bioscience, UK. Briefly, viral RNA was converted to cDNA then amplified using the ARTIC protocol v3 (LoCost) [25] and using V4 of the primer set, with sequencing libraries prepared using CoronaHiT as previously described [26]. Genome sequencing was performed using the Illumina NextSeq 500 platform (Illumina, CA, USA) with one positive control and one negative control per 94 samples. The raw reads were demultiplexed using bcl2fastq (v2.20). The reads were used to generate a consensus sequence using the ARTIC bioinformatic pipeline [27]. Briefly, the reads had adapters trimmed with TrimGalore [28] and were aligned to the WuhanHu-1 reference genome (accession MN908947.3) using BWA-MEM (v0.7.17) [29]. The ARTIC amplicons were trimmed, and a consensus was built using iVAR (v.1.3.1) [30]. PANGO lineages were assigned using Pangolin (v3.1.20) [31] and PangoLEARN model dated 2022-02-02 [32]. In this manuscript, we used the Pango lineage designation system.

## Acknowledgements

We thank the Lebanese Ministry of Public Health and the World Health Organization for supporting this work. We also thank the hospitals for their collaboration and facilitating samples storage and collection.

## Notes

### Competing Interest Statement

The authors have declared no competing interest.

